# The Achilles’ heel of coronaviruses: targeting the 5’ Polyuridines tract of the antigenome to inhibit Mouse coronavirus-induced cell death

**DOI:** 10.1101/2021.07.26.453908

**Authors:** Hemayet Ullah, Saadyah Averick, Qiyi Tang

**Author notes:** Corresponding author: Hemayet Ullah.

## Abstract

The current coronavirus pandemic situation is worsened by the rapidly-spreading SARS-CoV-2 virus variants. Identification of viral targets that are indispensable for the virus can be targeted to inhibit mutation-based new escape variant development. The 5’-polyU tract of the antigenome offers such a target. Host cells do not harbor 5’-polyU tracts on any of their transcripts, making the tract an attractive, virus-specific target. Inhibiting the 5’-polyU can limit the use of the tract as template to generate 3’ polyA tails of +RNAs of coronaviruses. Here, a modified DNA oligo with 3’ polyAs is used to target the 5-polyU tract in mouse coronavirus (MHV-A59). The oligo treatment in mouse 17CL-1 cells infected with MHV-A59 significantly prevented virus-induced cell deaths. This proof-of-concept result shows a unique mode of action against mouse coronavirus without affecting host cells, and can be used for the development of novel classes of drugs that inhibit coronavirus infection in host cells, specifically by the COVID-19-causing virus SARS-CoV-2. In addition, as the 5’-polyU tract is immediately generated upon infection, the tag can also be targeted for reliable early detection of viral infection.

## Introduction

The global havoc brought about by the COVID-19 pandemic is becoming more exacerbated by the rapidly spreading etiological SARS-CoV-2 virus variants [1]. Though vaccination against the virus has paced up, apprehensions surrounding so-called immune-escape variants raise concern for a worsening of the pandemic. Lack of enough vaccines have caused many countries to extend the gap between the two doses of vaccinations, which may lead to long durations of intermediate levels of immunity that could accelerate the emergence of new variants [2]. The potential consequences of emerging variants are increased transmissibility, increased pathogenicity and the ability to escape natural- or vaccine-induced immunity [3]. Drugs with high efficacy against the virus have not been developed yet. Concerns also surround whether drug-escape-variants, like immune-escape-variants, can evolve against any drug developed targeting the virus. Targeting coding regions directly can potentially lead to the evolution of escape variants.

In this regard, mitigation strategies should consider a mechanism to prevent the development of any escape-variants. An effective targeted strategy would be to block an area of the virus genome that is absolutely required by the virus to complete its life cycle. Any mutation in the targeted area will not be tolerated by the virus; thereby, the block will limit the virus’s ability to mutate into a new variant in the presence of selection pressure from drugs and/or vaccines. In this regard, the 5’ polyuridines (5’ poly-U) tract on the antigenome (minus RNA genome) of the positive RNA virus offers such a target (patent application)^1^. The 5’poly-U tract on the antigenome is indispensable for the virus as it is used as a template to generate the 3’ poly-A tail in its +RNA genome and the poly-A tail serves as template to generate the 5’ poly-U tract on the antigenome [4, 5]. Any mutation within the 5’poly-U tract will most likely prevent the virus from generating its 3’ poly-A tail, which would effectively hamper the generation of its full-length genome to complete its life cycle. The essentiality of the homopolymorphic 5’ poly-U tract on the antigenome for the survival and infectivity of positive RNA viruses, including many coronaviruses, is well-established [5–7]. On the other hand, host cells do not harbor any 5’poly-U tracts on any of their transcripts, making the tract an attractive, virus-specific target. Eukaryotic RNA polymerase III transcribes some small RNAs (t-RNA and 5S rRNA) with 3’ poly-U tract as a part of the transcription termination mechanism [8]. Molecular distinction between the viruses’ 5’ Poly-U tract from the host’s 3’ poly-U tract on transcripts will be a central consideration to avoid any off-target effects. Here, we successfully used a simple approach to this problem of targeting the 5’-polyU of a mouse coronavirus (MHV-A59) antigenome by providing a complementary 3’-polydAcontaining oligo. Use of the oligo against a fluorescent mouse coronavirus (MHV-A59) clearly indicates that the treatment of cells infected with the coronavirus are protected from the virus-induced cytopathic effects. This work represents a novel avenue of potential coronavirus treatment strategies, in its use of oligonucleotide-based therapies for coronaviruses infection that spare host cells.

## Results

### Design of DNA oligo complementary to the MHV-A59 5’ end of the antigenome (minus RNA genome)

The oligo sequences were carefully designed to be complementary to the polyUs at the 5’ end of the minus strand of the MHV-A59 genome (Fig 1A). The modifications were done to the oligos to provide increased resistance to nuclease degradation, reduced toxicity, and to provide increased affinity for binding to complementary RNA (Fig. 1B). Though these kinds of modification have been used in antisense technology to force usage of alternative polyadenylation site selection and to increase abundance of message [9], here the use of polyA oligo, rather than gene sequence targeted oligo, can minimize any such effect on the genome. Though there are 11 flanking bases running into the 3’ end of the N gene sequence of the virus, lack of any cryptic polyadenylation signal in the vicinity of the targeted sequence reduces any possibilities of use of alternative polyadenylation site.

**Fig.1:**
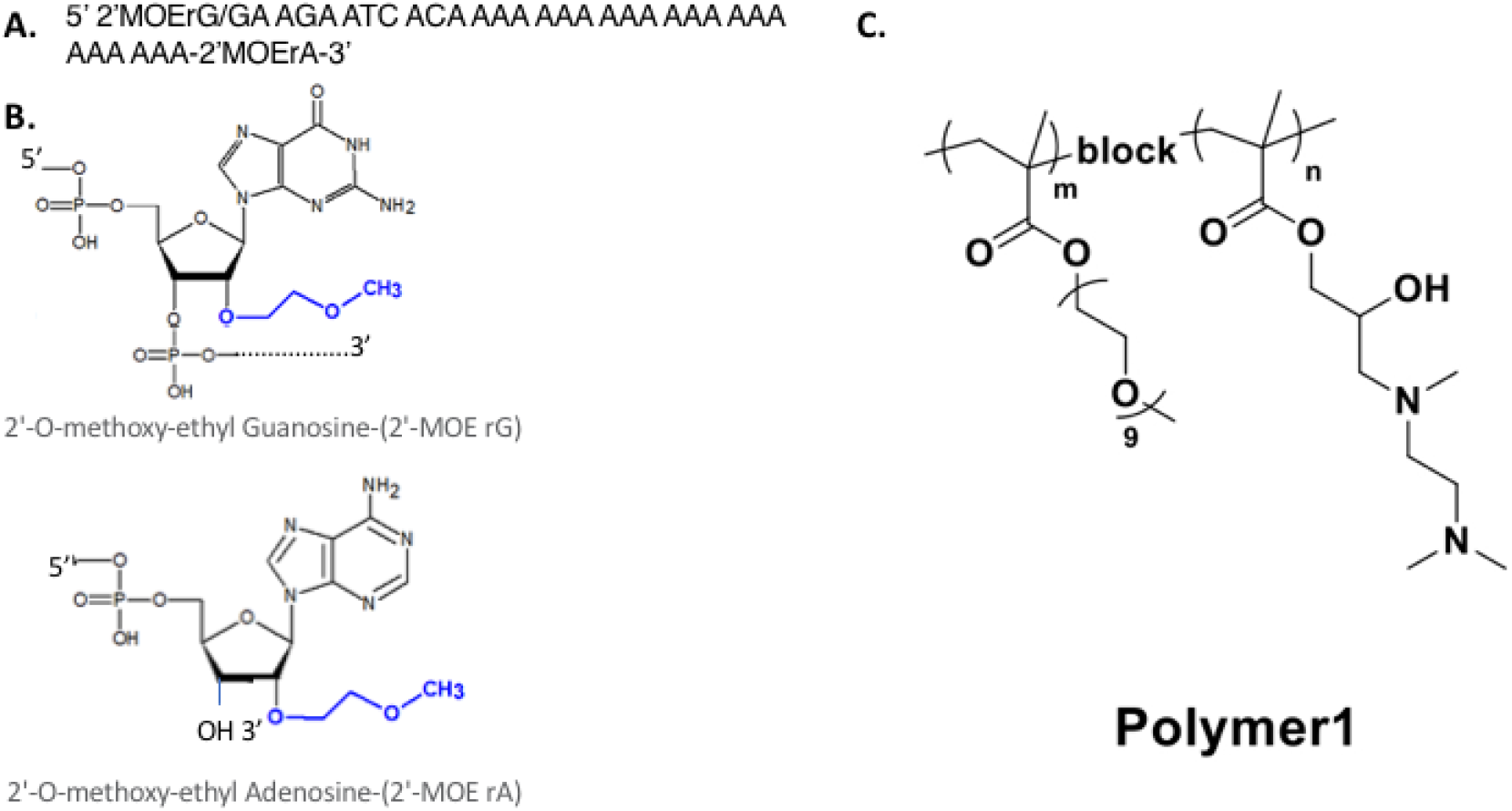
Sequence, modifications of DNA oligo used to target 5’-polyU of mouse coronavirus. A. The oligo sequence is designed to be complementary to the 5’ end of the minus strand of Murine hepatitis virus strain MHV-A59 genome (Accession number NC_001846 of whole genome is used to generate the complementary sequence). Note 11 bases complementary to sequences downstream of 5’polyU is added to allow hybridization to the 5’-polyU instead of any cellular 3’-polyU tags. B. 5’ and 3’ modifications are (2’-O-methoxy-ethyl bases). C. Structure of the block-copolymer used to deliver the oligos to the host cell.

### Oligo carrier nano-particles

Barriers to the successful translation of oligonucleotide-based therapeutics are the poor drug-like qualities of oligos. Oligonucleotides are relatively large (several kDa), anionically-charged macromolecules that are readily degraded by endogenous nucleases. These aforementioned properties of oligonucleotides prevent their cellular penetration, limit circulatory half-life, and have potential immunogenic effects. To overcome these challenges, efforts have been made to create of drug delivery vehicles capable of complexing oligonucleotides to neutralize their charge and prevent nuclease-based degradation. These vehicles are typically comprised of a quaternary or tertiary amine cationic moiety, other heteroatoms are possible, attached weight to a polymer backbone or lipid. Each drug delivery has distinct advantages and disadvantages including synthetic accessibility, oligonucleotide:carrier stability, pharmacokinetic half-life, ease of targeting agent inclusion, among others. Through judicious optimization and tuning of the structure of the drug delivery vehicle, including the addition of targeting agents, the clinical potential of oligonucleotide-based therapeutics has achieved initial success. Oligonucleotide delivery: To determine if the delivery vehicle would impact the efficacy of the Gen 2 DNA oligonucleotide, the oligo was formulated with a lipid-based delivery agent (lipofectamine) or a block-copolymer. The oligonucleotide was complexed via electrostatic interactions of the negatively-charged DNA backbone and cationic charge on the lipid/polymer systems. The lipid-based oligonucleotide delivery systems are currently employed in the mRNA based delivery of SARS-Cov2 vaccines, albeit with unknown lipid structures. The Polymer1 system, prepared as previously described is a block copolymer with a first block that provides a hydrophilic biocompatible shell of oligoethylene oxide methacrylate and second block comprised of a hydrophilic tertiary amine methacrylate [10]. The design of this system allows for complexation of the oligonucleotide while protecting from nucleases and non-specific binding. These two divergent delivery vehicles were tested to ensure that the observed effects of oligonucleotide transfection were not carrier dependent and to screen for potential in vivo oligonucleotide carriers. However, here we report the results from Polymer1-based transfection.

### Oligos prevent virus-induced cell death

Mouse Coronavirus-MHV-A59 was obtained from the NIH/NIAID BEI Resources stock center (Cat# NR-53716). The stock is a virus preparation made using targeted recombination and selection for MHV with stable and efficient expression of the gene encoding eGFP. The eGFP gene was inserted into the MHV genome in place of the nonessential gene ORF4 in mildly neurovirulent strain, MHV-A59 [11]. Coronaviruses are known to develop double membrane vesicles (DMVs) by modifying cytoplasmic membranes in the perinuclear area to anchor the replication and transcription complex in order to support viral replication and RNA synthesis [12]. For coronaviruses, replicase nonstructural protein 4 (nsp4) has been proposed to function in the formation and organization of replication complexes [12]. We used an excess concentration of oligos to ensure that enough oligos will be available for the cytoplasm-localized Replication-Transcription complex (RTC), which the virus uses to develop its genomes and subgenomic RNAs. Serial dilution of the stock virus was used to infect the mouse fibroblast cell line 17CL-1 that was derived by spontaneous transformation of 3T3 cells. The cells were maintained on MEM medium supplemented with 10% FBS. In order to investigate the effect of the designed oligos on the mouse coronavirus, almost 90% confluent 17CL-1 cells were infected with the MHV-A59-eGFP at 0.01 MOI. After one hour of adsorption, the media were removed and the cells were washed three times with PBS buffer. Four μg of oligos were thoroughly mixed with 4 μg of polymer1 and was incubated at room temperature for 15 minutes before adding to the virus infected cells. As a control, 4 μg of Polymer1 only was added to the virus infected cells. Bright field images were taken after 48 hours of incubation at 37_0_C in a 5% CO2 incubator. As can be seen from the Fig.2, the cells treated with only Polymer1, the virus was able to cause significant cell deaths as the monolayer detached cells appeared as round, fused, aggregated. Not many intact cells were seen in the Polymer1 only treated cells. The similar effect is significantly attenuated and delayed in the oligo treated cells (Fig.2B). Significant numbers of cells protected from virus cytopathic effects were seen in the oligo treated culture wells. As the cells were treated with the virus before the oligo treatment, the effect may not be emanated from the exclusion of viruses from the cells. Though the binding assay of the oligos to the target sequences is still pending, the observed results strongly implicate the oligos in the protection of the virus-infected cells.

**Fig.2:**
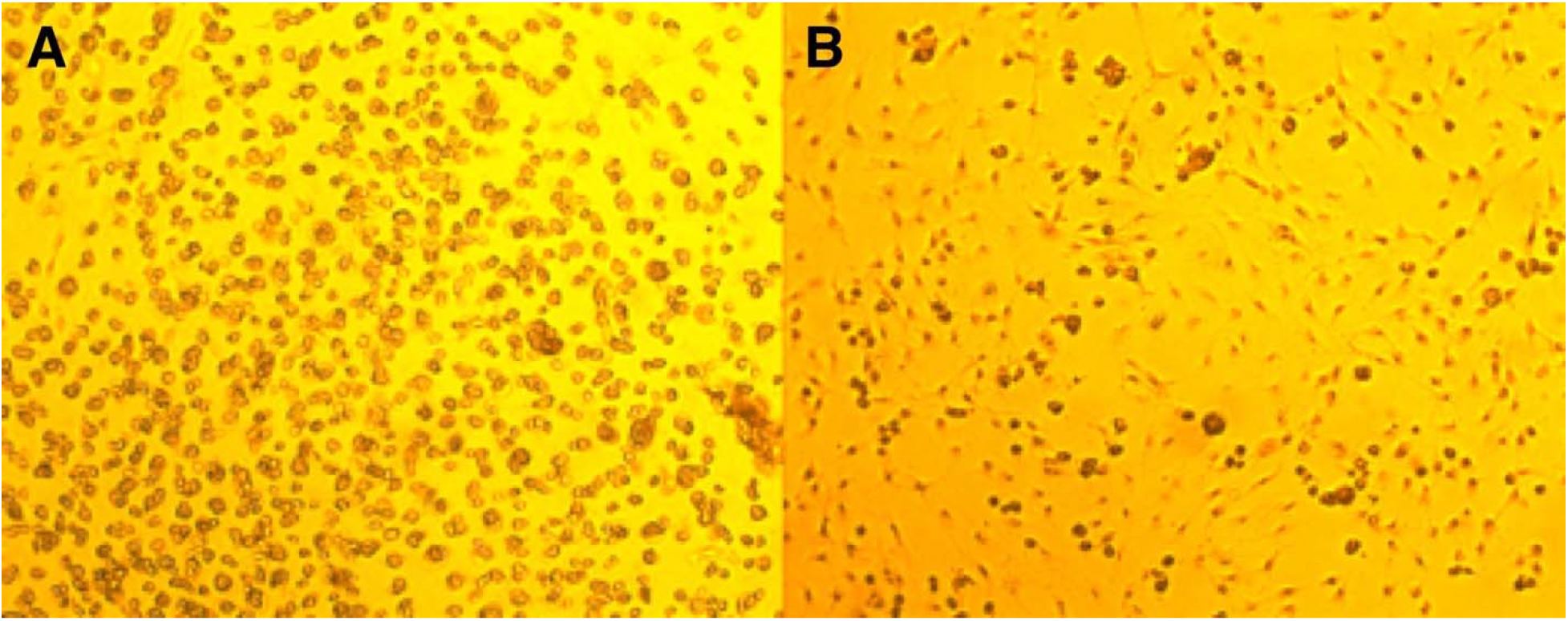
The DNA oligos prevent MHV-A59 induced death of mouse 17CL-1 cells. The image was taken 72h after infection. Image represents infection with virus at 0.1 MOI. A. MHV-A59 with Polymer1 only. B. MHV-A59 with Polymer1 and oligos. Oligos were added after one hour of adsorption of the virus. Rounded, aggregated, fused, and granulated cells rapidly detaching from the monolayer are shown in panel A. (Original magnification 10X).

We also investigated the virus proliferation by reading out GFP expression 24h post-infection. Fig. 3 shows that in the oligo-treated cells (Fig. 3B), the GFP expression is not as widespread as that in cells that did not receive oligo treatment (Fig. 3A). The oligos were added post-adsorption of the viruses. Though GFP expression is seen almost all the cells in the frame (Fig. 3A), the oligo-treated cells did not show as intense and widespread expression 24h post infection. Though the replication of the viruses is not significantly inhibited by the oligos treatment, the proliferated viruses were not able to induce cell death as profoundly as the viruses without the oligo treatment (Fig. 2).

**Fig. 3:**
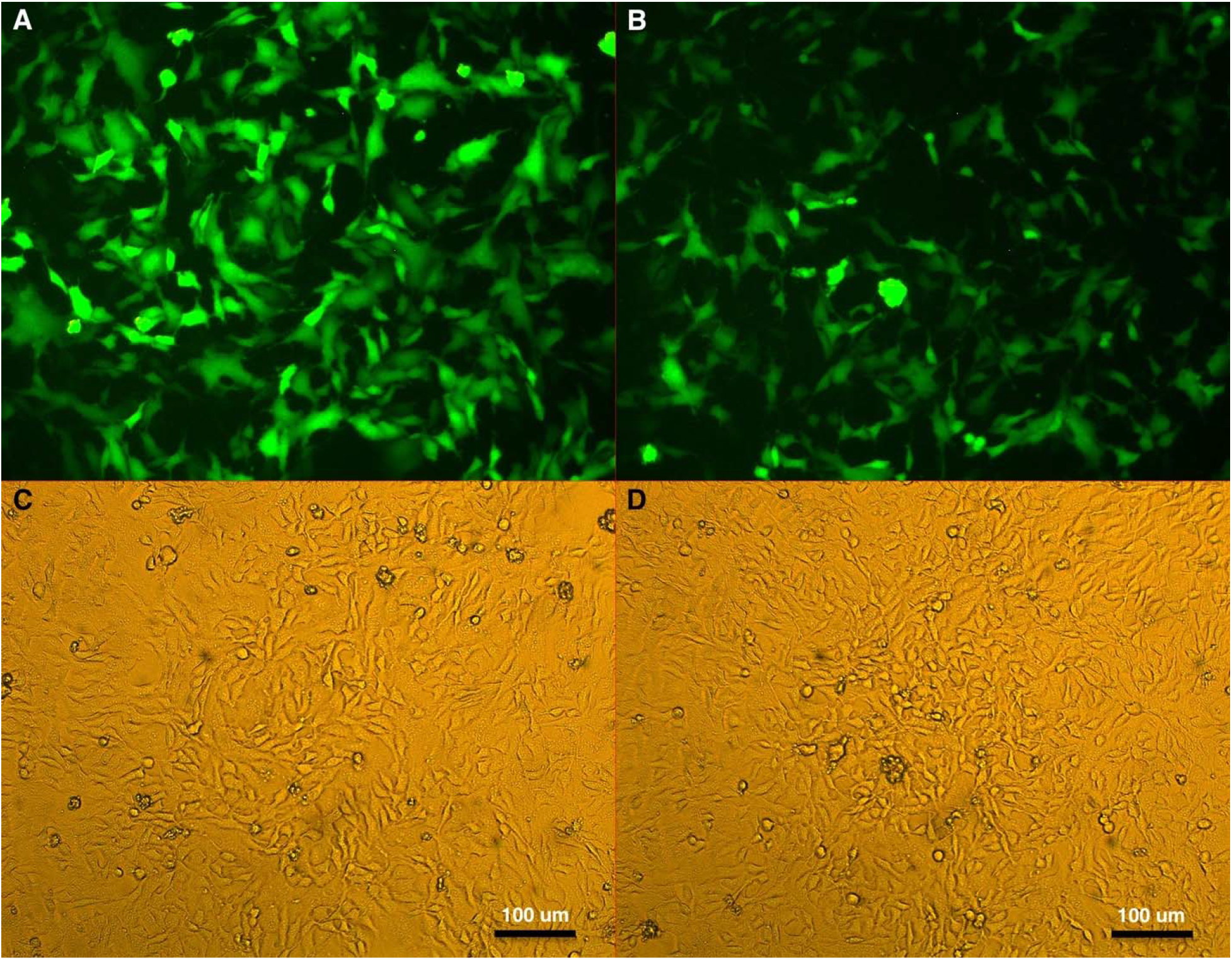
Mouse 17CL-1 cells treated with MOI 0.1 MHV-A59-eGFP and imaged 24h post-infection. A. Fluorescence image of Polymer1-only treated cells; B. Same frame as in panel A with bright field image. C. Fluorescence image of cells treated with oligos; D. Same frame as in panel B with bright field image. Scale bar: 100 μm.

In order to investigate virus induced-cell death, we fixed the cells 72h post-infection with 4% paraformaldehyde (PFA) in PBS and stained the cells with 0.5% crystal violet in 25% methanol. Before fixing the cells, the floating cells and cell debris were removed and mostly the cells remaining in the monolayer were fixed and stained. As can be seen in Fig.4, the oligo-treated cells showed more intact cells (Fig. 4B) whereas the only Polymer1 treated cells had only few cells remaining in the monolayer (Fig. 4A).

**Fig. 4:**
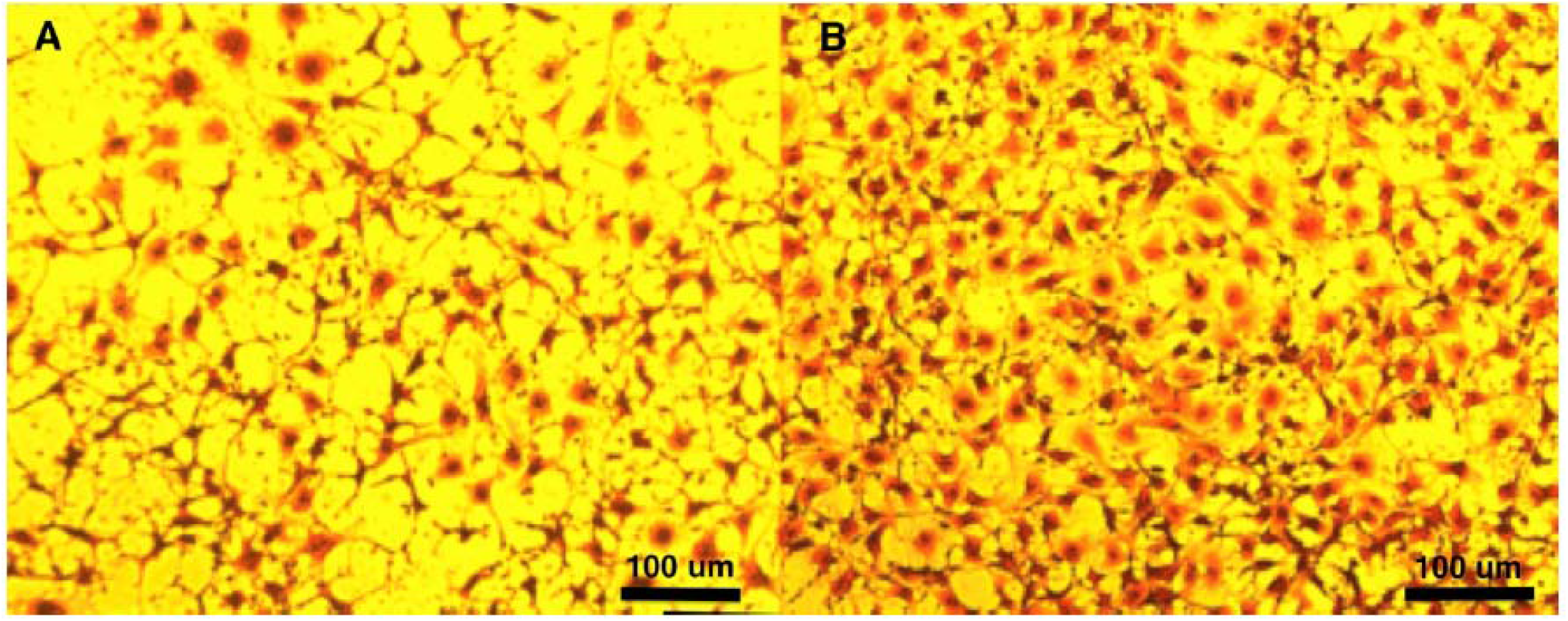
PFA fixed and crystal violet stained 17CL-1 cells 72h post virus infection at MOI 0.1. Original magnification-10X. Here the cells were pre-treated with oligos for 1h followed by 1h of virus infection. Scale bar: 100 μm.

We also used 0.4% trypan blue-based live and dead cell assays to investigate the total number of live and dead cells 48h post infection (Table I). As can be seen from Table I, while more than 93% of the cells were viable with the oligo treatment, only 76% of the cells were viable without the oligo treatment. In order to eliminate any effect from the oligos on the survivability of the cells, no virus-treated cells were also counted. In the absence of any virus treatment, we found no significant differences in cell survival in the Polymer1 alone or in oligo-treated cells.

**Table I:** Oligo treated cells have more viable cells after 48 hours of virus infection

| Treatment | MHV-A59 | #of cells/ml | % of live cells (±S.E.) |
| --- | --- | --- | --- |
| Polymer1 (4 μg) | No Virus | $5.95 \times 10^6$ | $91.8 \pm 0.99$ |
| Polymer1 + Oligos (4 μg each) | No Virus | $6.46 \times 10^6$ | $93.4 \pm 1.13$ |
| Polymer1 (4 μg) | Yes | $1.02 \times 10^7$ | $76.0 \pm 2.89$ |
| Polymer1+ oligos (4 μg each) | Yes | $9.68 \times 10^6$ | $93.3 \pm 1.45$ |

Cells from three different wells were counted for each treatment. S.E.: Standard Error

As the total cell counts and number of live cells in the absence of the virus are not significantly different in the Polymer1 alone or Polymer1 with oligo-treated cells (Fig. 5), it is unlikely that the oligos themselves prevent cell death; rather the oligos prevent virus induced-cell death. Fig. 5 shows that the cells in the absence of virus but in the presence of polymer1 alone or polymer1 with oligos have essentially same confluency. The abundance of virus-induced cell clumps as seen in Fig. 2 and the spread of virus through cell-cell fusion as seen in GFP fluorescence in Fig. 3 indicate that the oligos can be used as a therapy option for preventing virus-induced host cell death.

**Fig. 5:**
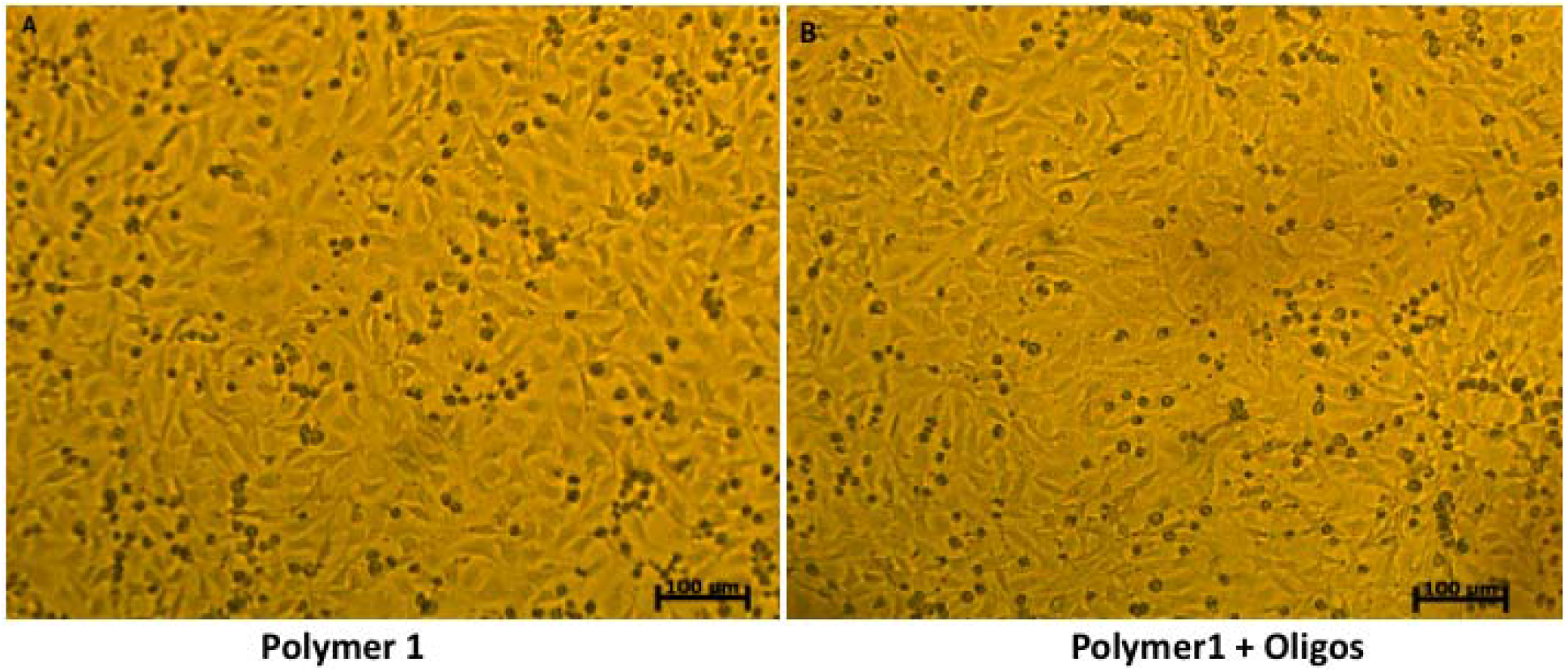
Bright field images of 17CL-1 cells without any virus treated with either Polymer1 alone (A) or with Polymer1 and oligos (B).

## Discussion

A powerful way to contain the current pandemic of COVID-19 is to target the virus in such a way that the virus is unable to generate escape variants by mutating its genome sequences. In this regard, we propose to target the 5’-polyU tag on its negative RNA strand so that the virus is unable to use the tag as a template to generate the 3’ polyA tails needed for its efficient translation using host translation machinery. Any mutation on the polyU tag will is unlikely to be tolerated by the virus, as this homopolymorphic stretch of polyUs are used as a template to generate the polyA tails. Moreover, as the host cells do not maintain any transcripts with 5’-polyU tags, the target offers a virus-specific, unique tag that can be targeted without affecting any host cell transcripts. Here as a proof-of-concept, we targeted the mouse coronavirus 5’-polyU tag with a simple DNA oligo containing polydA that was designed based on the published genome of the mouse coronavirus. We show that the oligo approach is useful as the oligo was successful in preventing and delaying virus-induced host cells death. Though the oligos were not able to completely inhibit virus replication, the prevention/delaying of cell death can be used as evidence that the oligos binding to the PolyU may not prevent replication, but rather may prevent synthesis of fully functional virions. Further experiments with infectivity potential of the oligo-treated viruses may shed more mechanistic insights into the target. Here we have presented preliminary evidence to establish the 5’-polyU tag as a legitimate target for the successful containment of virus spread.

The successful application of the oligo to target the mouse coronavirus 5’-polyU tag can be attributed to the unique design of the oligo and to the use of block-polymers to deliver the oligos to the cell interior. The coronaviruses never enter the host cell nucleus where host cell mRNAs are processed by nuclear localized enzymes, the coronaviruses are known to replicate in the cytoplasm [13]. This may allow the oligos to bind to their target easily.

In a recent paper on the betacoronavirus mouse hepatitis virus strain A59 (MHV-A59) and on the alpha CoV enteropathogenic porcine epidemic diarrhea virus (PEDV), it was shown that virus endoribonuclease (EndoU) facilitates evasion of host pattern recognition receptor MDA5 by cleaving the 5’-polyuridines from the viral minus RNA strand which is a product of the 3’ poly-A-templated RNA synthesis [14]. The paper suggested targeting the EndoU to allow the MDA5 to mount a robust interferon-based immune response. Though this result suggests that coronaviruses shorten the 5’poly-U tract to evade the host’s interferon-based immunity, shortened poly-U tracts are always found to be maintained at the 5’ end of the minus RNA strand, indicating the essential need for the tract. The role of the 5’poly-U tract in completing the coronavirus’s life-cycle is well described [5, 6, 14, 15]. In contrast to cellular mRNAs, in which the poly(A) is non-templated and is added post-transcriptionally, the viral poly(A) is template coded from a 5’ poly-U tract in positive RNA viruses like picornavirus [16] and in coronaviruses [15, 17]. Even though N-terminal RdRp-associated nucleotidyl-transferase activity can potentially add template-independent poly A, in HCoV-228E-a beta-coronavirus, it has been found that nsp8 (viral RdRp accessory subunit) has efficient 3’-terminal adenylyltransferase (TATase) activity only when the other strand has the 5’ poly-U tract[18]. The oligo(U)-assisted/templated TATase activity on partial-duplex RNAs was confirmed for two other coronavirus nsp8 proteins, suggesting that the activity is conserved among coronaviruses [18]. The immature positive RNA strand becomes mature and stabilized only when iterative replication from the poly-U template adds the 3’ poly-A tail [6]. It is quite possible that binding of the oligos to the 5’-polyU tag may mimic a shortened 5’-polyU stretch which can allow short term evasion of the host immunity response, however, this may prevent generation of the fully functional and virulent virions with full potential of infection capabilities. More mechanistic experiments will be able to establish the molecular details of such reduced capabilities to cause cell death as reported in this study.

The above discussion indicates that the positive sense RNA viruses essentially require a 5’ poly-U tract to complete their life cycle and to successfully infect host cells. Therefore, inhibiting/blocking the 5’ poly-U tract should also interfere with this key survival mechanism of the virus and may lead to the production of a ‘dead’ genome or to a crippled virus unable to infect. Targeting the 5’ poly-U with lipid nanoparticle wrapped poly-A or with oligo dA, or DNA aptamer, or 5’ Poly-U binding small compound and/or protein can limit the use of the minus strand for generating the positive sense RNA strand genome and the subgenomic mRNAs with poly-A tails.

In addition, it has not escaped our attention that the poly-U tract is immediately generated upon infection [15]. As such, the tag can be leveraged and targeted for early detection of viral infection as well. This may overcome many of the issues encountered with the prevalently used real-time PCR-based detection systems, such as the lack of enough template early in the infection which may drive false negative results.

Scientists around the world are working to understand this devastating illness and help put an end to the terrible toll caused by the COVID-19 pandemic. Experimental establishment of an indispensable target like this 5’poly-U tract on the antigenome of coronaviruses may assist in the development of effective drugs that prevent the generation of drug-escaping virus variants.

## Acknowledgement

The following reagent was obtained through BEI Resources, NIAID, NIH: Recombinant Murine Coronavirus MHV-A59 with Enhanced Green Fluorescent Protein (eGFP), NR-53716. The following reagent was obtained through BEI Resources, NIAID, NIH: Murine 17Cl-1 Cell Line (derived from 3T3 cells), NR-53719.

## Footnotes

1 Provisional patent application # 63025613, EFS ID# 39456605 (Title: Targeting the Unique minus Strand mRNA to Stop Replication of COVID-19 Causing SARS-CoV-2 virus without affecting host cells). PCT application submitted May 14, 2021.

